# The joint associations of physical activity, sedentary time, and sleep on VO_2max_ in trained and untrained children and adolescents: A novel five-part compositional analysis

**DOI:** 10.1101/2022.09.21.508871

**Authors:** A. Runacres, K.A. Mackintosh, S. Chastin, M.A. McNarry

## Abstract

The benefits of physical activity (PA) and the negative impacts of sedentary time (SED) on both short- and long-term health in youth are well established. However, uncertainty remains about how PA and SED jointly influence maximal oxygen uptake (VO_2max_). Therefore, the aim of this study was to determine the joint influence of PA and SED on peak VO_2_ using compositional analyses. 176 adolescents (84 girls, 13.8 ± 1.8 years) completed an incremental ramp test, and supramaximal validation bout, on cycle ergometer with PA and SED recorded for seven consecutive days on the right hip using a ActiGraph GT3X accelerometer. Time spent in sleep, SED, light, moderate and vigorous PA was analysed using a compositional linear regression model. Compositions with 10 minutes more time in vigorous PA (> 27.5 mins·day^-1^) compared to the average 17.5 mins·day^-1^ were associated with a + 2.9% - 11.1% higher absolute and allometrically scaled peak VO_2_. Whereas compositions with less (> −10 mins·day^-1^) VPA were associated with a reduced absolute and allometrically scaled VO_2max_ (−4.6% - 24.4%). All associations were irrespective of sex, maturity, and training status. The proportion of time spent sedentary had little impact on absolute and scaled peak VO_2_ (0.01 – 1.98%). These findings therefore highlight that intensity of PA may be of greater importance for increases in VO_2max_ than reductions in SED and should be incorporated into future intervention designs.

## 1. Introduction

Poor maximal oxygen uptake (VO_2max_) has been associated with an increased risk of cardiovascular and metabolic disease, leading to an increased likelihood of premature mortality across the lifespan [1–4]. VO_2max_, defined as the highest rate of oxygen consumption despite further increases in work rate [5], is also key to athletic performance, with youth athletes consistently reported to have a greater peak VO_2_ than their untrained counterparts [6–9]. Whilst training is well-established to improve VO_2max_ in youth athletes [6,9–11], what remains less clear is the influence of habitual physical activity (PA) and sedentary time (SED) on VO_2max_. More specifically, some studies have reported a significant association between PA and VO_2max_ [12–14], whereas others argue that children and adolescents rarely experience PA of a sufficient duration and intensity to significantly influence VO_2max_ [11,15].

Moderate-to-vigorous PA (MVPA) is perhaps the most widely used PA metric in children and adolescents [16–18] and remains the focus of governmental physical activity guidelines [19,20]. Physical activity research in children has consistently suggested that increased levels of MVPA is positively associated with VO_2max_ in youth [12–14], and excess SED is negatively correlated with VO_2max_ [21,22]. However, what remains unclear is the joint association between VPA and SED with elite junior athletes reporting similar, if not more, SED than their inactive, sedentary peers [23]. Therefore, whether VPA can offset the negative effects of SED requires urgent investigation and cannot be done with the reliance on correlational statistics, and the use of linear regressions modelling which assume independence between variables [23,24]. Indeed, traditional correlational statistics are inappropriate to account for the constrained and co-dependent nature of PA data, potentially creating spurious associations.

Compositional analysis allows all movement behaviours to be expressed as a proportion of a finite period, enabling the individual, and joint, effects of movement behaviours on outcome variables to be established [16–18,25,26]. Consequently, compositional analysis could provide novel insights into the joint influence of movement behaviour, intensity, and volume on VO_2max_. Indeed, Carson et al. [16] found that the overall movement behaviour composition explained ~38% of the variance in the VO_2max_ of 4,169 Canadian children and adolescents (8 – 17 years). Despite this, when 10 minutes of time was allocated to, or removed from, MVPA, there was a negligible effect on VO_2max_, with predicted changes ranging from 0.03% - 0.05% [16]. This could be due, at least in part, to some methodological inconsistencies including the estimation of peak VO_2_ from a field-based test which is likely to misrepresent true cardiorespiratory fitness [27], the pooling of data from boys and girls despite the well-established physiological differences [28], and the failure to account for maturity or training status. Alternatively, Carson et al [16] only explored the effect of changing MVPA compositions thereby combining MPA and VPA, which may potentially mask the importance of the intensity of physical activity for improving VO_2max_ in youth. Indeed, training studies consistently show that significant improvements in absolute, and allometrically scaled, VO_2max_ only occur when the intensity is sufficiently vigorous [10,29]. Indeed, Gutin et al. [30] reported a stronger association between VPA and VO_2max_ (r^2^ = 0.43, p < 0.01) than MPA (r^2^ = 0.30, p < 0.01) in adolescents and these findings have been corroborated by both Dencker et al. [12] and Latt et al. [14] who reported that the amount of time spent in VPA explained 9.0 – 15.8% of the variance in peak VO_2_ in children and adolescents.

Therefore, the aim of this study was to examine the independent, and interactive, effects of the five movement behaviours (SED, LPA, MPA, VPA and sleep) on VO_2max_ in children and adolescents. The second aim was to explore the effect of baseline fitness, sex, and maturity on the predicted changes in VO_2max_ elicited by changing PA compositions.

## 2. Methods

Ethics approval was granted by the institutional research ethics committee at Swansea University prior to the commencement of data collection and the study conformed to the Declaration of Helsinki. Before participants were accepted into this cross-sectional study, written informed parental consent and participant assent were obtained, along with all parents completing a pre-screening medical questionnaire on behalf of their child. Participants were excluded if they had known cardiovascular, metabolic, kidney, or any other disease that meant they would not have been able to complete the exercise protocol. The trained children and adolescents were all national level athletes who were part of a long-term athlete development (LTAD) program overseen by the national governing body (NGB) of their sport (Hockey, Football and Gymnastics). Untrained participants were recruited from local schools across South Wales and were not formally engaged in sport training outside of curricular physical education lessons. The final sample consisted of 237 participants encompassing 108 trained (43 girls; age: 13.5 ± 2.1 years) and 129 untrained (51 girls; 13.8 ± 1.4 years) children and adolescents.

### 2.1 Experimental Procedures

All participants were required to attend one session at which they initially had their stature and sitting height measured to the nearest 0.1 cm using a Holtain Stadiometer (Holtain, Crymych, Dyfed, UK) and their body mass measured to the nearest 0.1 kg using electronic scales (Seca 803, Seca, Chino, CA, USA). Maturity status was subsequently estimated using the equations of Mirwald et al. [31], with participants deemed pre-pubertal, pubertal, and post-pubertal if they were more than one year from, within one year of, or more than one year post peak height velocity (PHV), respectively.

VO_2max_ was assessed using an incremental ramp test to volitional exhaustion on a cycle ergometer (Lode Excalibur Sport, Groningen, Netherlands) which started with a three-minute warm-up at 10 W before increasing by 20 – 25 W·min^-1^, depending on the participant’s age. All participants were instructed to maintain a cadence of 60 – 80 revolutions per minute (rpm) throughout the test, with volitional exhaustion defined as when participants could not maintain a cadence above 50 rpm. Inspired and expired air were measured on a breath-by-breath basis throughout the incremental ramp test using a Vyntus metabolic cart (VYAIRE medical Ltd, Mettawa, IL, USA). Following five minutes active and ten minutes passive rest, a supramaximal validation bout was performed [32]. Specifically, participants warmed up for a further three minutes at 10 W before a step-transition to 105% of the peak power achieved during the incremental ramp test. Participants were instructed to maintain a cadence above 50 rpm for as long as possible, with gas exchange measured continuously on a breath-by-breath basis throughout the exercise bout.

Participant’s habitual physical activity was subsequently measured for seven consecutive days using a ActiGraph GT3X (ActiGraph, Pensacola, Florida, USA) worn on the right hip, sampling at 100 Hz. Children and adolescents also completed a seven-day log to detail periods when the monitor was removed, waking time and time going to bed, to minimise the misclassification of non-wear time as sedentary time or sleep.

### 2.2 Data Analyses

The raw breath-by-breath VO_2_ data from both the VO_2max_ and supramaximal bout were averaged into 10-second bins, with the VO_2max_ defined as the highest 10-second moving average during the ramp incremental test. To aid comparisons between sex, maturity, and training sub-groups, VO_2max_ was allometrically scaled to account for body mass differences between participants [6,32]. Evenson et al. [33] cut-points were utilised to determine the time spent in each PA intensity which have been shown to be the most reliable for children and adolescents [34]. Sleep time and efficiency was calculated using the algorithms of Sadeh et al. [35]. Wear-time criteria was set as ≥ 8 hours on any three days. Using the Evenson et al. [33] cut-points, sleep algorithms, and wear-time each day was expressed as a five-part movement composition (SED, LPA, MPA, VPA, Sleep) and linear predictive models were employed to predict changes in VO_2max_. The smallest worthwhile change (SWC) in VO_2max_ (l·min^-1^) and allometrically scaled peak VO_2_ (ml·kg^-b^·min^-1^) was calculated for each sex, maturity and training sub-group using the formula 0.2*group SD [36]. The SWC was then subsequently presented as a percentage of the group mean to aid comparisons between all sub-groups.

All compositional analyses were conducted in R (http://cran.r-project.org) using the compositions package (version 1.40-2) and its dependencies [25]. Compositional geometric means were computed to indicate the proportion of time spent in each PA behaviour or sleep each day, by expressing each behaviour, after normalisation, as a proportion of the total time [18,25]. Variance matrices were calculated to provide an indication as to the dispersion and co-dependency of movement behaviours and were calculated by measuring the variance between pair-wise log ratios [25,26]. Specifically, a ratio tending towards zero indicates high co-dependency, with the numbers further from zero indicating less co-dependency. Sequential linear regression models were created by rotating each of the five behaviours via isometric log ratio (ILR) transformations to examine the relative effect of all movement behaviours on the VO_2max_ and allometrically scaled VO_2max_ [18,25]. The first coefficient and its *p* value were reported for each rotation to determine whether the individual movement behaviour was associated with the outcome variable relative to the other movement behaviours, and its relative significance. Additionally, the overall model significance (*p* value) and R^2^ value were reported to gain an insight into the variance explained by the overall movement composition. All movement behaviours were also sequentially mapped against each other, producing ternary heat maps displaying the predicted absolute and allometrically scaled VO_2max_ for each sex, training, and maturity group. Finally, change matrices were conducted to predict the change in absolute and scaled VO_2max_ by systematically reallocating 10 minutes from one movement behaviour to another [18,25,26]. All predictive changes were presented as a percentage change relative to the compositional mean, with significant changes identified as any change greater than the SWC (%).

### 2.3 Statistical analyses

All traditional statistical analyses were conducted in SPSS version 26 (IBM, Portsmouth, UK), with significance accepted as p < 0.05. Between group differences in anthropometric characteristics and absolute and allometrically scaled VO_2max_ were assessed using a MANOVA, with post-hoc tests with Bonferroni correction applied to identify the specific location of significant differences as appropriate.

## 3. Results

Of the original 237 participants, 61 were excluded for failing to meet the wear-time criteria, therefore 84 trained (40 girls) and 92 untrained (44 girls) children and adolescents were included in the final analyses. There were no significant differences in the anthropometrics of those included and excluded (p > 0.05). Post-pubertal adolescents were significantly older, taller, heavier, and more mature than the pubertal or pre-pubertal children (p < 0.01), with significant differences in the same parameters also evident between pubertal adolescents and pre-pubertal children (p < 0.05, Table 1). The trained children and adolescents were taller (*F*_(1,175)_ = 12.7, p < 0.01) and had a higher VO_2max_ (l·min^-1^) than their untrained counterparts (*F*_(1, 175)_ = 15.3, p < 0.01), which persisted even after allometric scaling (*F*_(1,175)_ = 18.7, p < 0.01). Overall, boys had a higher absolute and allometrically scaled VO_2max_ than their female counterparts, irrespective of training and maturity status (*F*_(1,175)_ = 19.7, p < 0.01). VO_2max_ increased with maturity, irrespective of sex and training status (*F*_(1,175)_ = 16.2, p < 0.01), but there was no significant difference between any maturity group for allometrically scaled VO_2max_. There were no significant training, sex, or maturity interactions for any anthropometric variable or VO_2max_, regardless of how it was expressed.

**Table 1.**
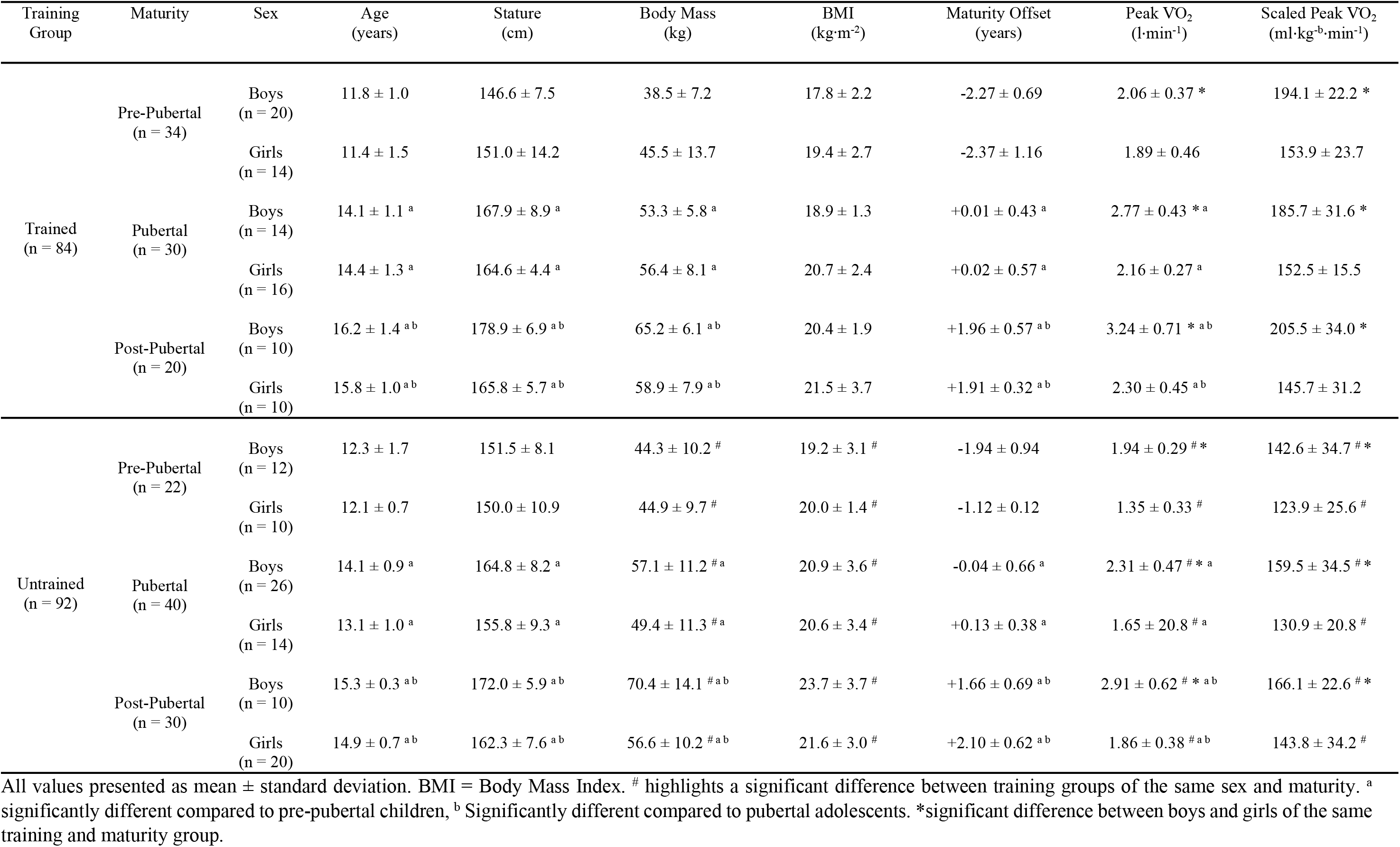
Participant descriptives.

In the trained participants, the geometric means highlight that the biggest portion of the day was spent in SED (41.2%), followed by sleep (39.2%), with VPA only accounting for 1.6% of the day (Table 2). Similarly, untrained children spent the longest periods of the day in SED (44.1%) and sleep (40.0%), with VPA making up just 1.3% of the day. Trained athletes completed more LPA (*F*_(1,175)_ = 38.1, p < 0.01) and VPA (*F*_(1,175)_ = 18.6, p < 0.01), but spent significantly less time sleep (*F*_(1,175)_ = 3.8, p = 0.05) compared to untrained participants, irrespective of sex or maturity. LPA and sleep, and SED and LPA, demonstrated the smallest variation and therefore high co-dependency, whereas VPA had the largest pair-wise log ratio variances compared to all other PA behaviours, indicating less co-dependency (Table 3). The ILR model revealed that the composition of PA, SED and sleep significantly predict both VO_2max_ and allometrically scaled VO_2max_ (Table 4). Additionally, the overall movement composition explained 48.7% and 37.7% of the variance in absolute, and allometrically scaled, VO_2max_, respectively. In isolation, when compared against all other PA intensities, only VPA significantly predicted allometrically scaled VO_2max_ (Y_VPA_ = 6.91, p < 0.02), with no significant individual associations evident for absolute VO_2max_.

**Table 2.** Geometric Means for the whole sample.

| Trained Athletes |  |  |  |
| --- | --- | --- | --- |
|  | Overall Mean<br>(minutes·day<sup>-1</sup>) | Geometric Mean<br>(minutes·day<sup>-1</sup>) | % of 24 hours |
| SED | 515.6 | 594.4 | 41.2 |
| LPA | 186.4 | 214.9 | 14.9 |
| MPA | 37.4 | 43.1 | 3.0 |
| VPA | 19.9 | 22.9 | 1.6 |
| Sleep | 489.7 | 564.5 | 39.2 |
| Untrained Controls |  |  |  |
| SED | 517.2 * | 634.5 * | 44.1 * |
| LPA | 134.4 | 164.9 | 11.5 |
| MPA | 37.9 | 46.5 | 3.2 |
| VPA | 15.2 * | 18.6 * | 1.3 * |
| Sleep | 469.7 | 575.5 | 40.0 |

SED = Sedentary Time, LPA = Light Physical Activity, MPA = Moderate Physical Activity, VPA = Vigorous Physical Activity, \*Indicates significant difference between training groups

**Table 3.** Pair-wise log ratio variation matrix in the full sample.

|  | SED | LPA | MPA | VPA | Sleep |
| --- | --- | --- | --- | --- | --- |
| SED | - | -0.018 | -0.029 | -0.053 | 0.023 |
| LPA | -0.018 | - | -0.039 | -0.042 | -0.016 |
| MPA | -0.029 | -0.039 | - | -0.020 | -0.019 |
| VPA | -0.053 | -0.042 | 0.020 | - | -0.045 |
| Sleep | 0.023 | -0.016 | -0.019 | -0.045 | - |

SED = Sedentary Time, LPA = Light Physical Activity, MPA = Moderate Physical Activity, VPA = Vigorous Physical Activity

**Table 4.** Model ILR parameters for Peak VO_2_ and scaled peak VO_2_.

|  | Model p<br>value | Model<br>R<sup>2</sup> | Y<sub>SED</sub> | p | Y<sub>LPA</sub> | p | Y<sub>MPA</sub> | p | Y<sub>VPA</sub> | p | Y<sub>Sleep</sub> | p |
| --- | --- | --- | --- | --- | --- | --- | --- | --- | --- | --- | --- | --- |
| Peak VO<sub>2</sub> (l·min<sup>-1</sup>) | < 0.001 * | 0.486 | < 0.001 | 0.993 | -0.070 | 0.112 | 0.041 | 0.483 | 0.029 | 0.529 | < 0.001 | 0.997 |
| Scaled peak VO<sub>2</sub><br>(ml·kg<sup>-b</sup>·min<sup>-1</sup>) | < 0.001 * | 0.377 | -0.627 | 0.865 | 2.390 | 0.396 | -5.606 | 0.136 | 6.914 | 0.019* | 1.709 | 0.668 |

All models were covaried for training status, sex and maturity. \* indicates a significant predictor of outcome variable. SED = Sedentary Time, LPA = Light Physical Activity, MPA = Moderate Physical Activity, VPA = Vigorous Physical Activity.

VPA was the most influential PA behaviour for absolute, and allometrically scaled, VO_2max_, irrespective of training status and sex, but the influence of PA behaviours was less clear in pubertal and post-pubertal adolescents (S1 Figures 1 and 2). Compositions with 10 minutes difference in any movement behaviour (SED, LPA, MPA, VPA, Sleep) had a minimal effect on peak VO_2_ in trained children and adolescents (Table 5), with all changes smaller than the SWC (S1 Table 1). Contrastingly, in pre-pubertal untrained children, compositions with 10 minutes less VPA (−5.2 mins·day^-1^) compared to the average 15.2 mins·day^-1^ was associated with a decrease in peak VO_2_ of between 3.2 – 10.5% (Table 6). Additionally, compositions with less LPA, but more MPA or VPA was associated with an increase in peak VO_2_ between 5.2% and 5.8% in pre-pubertal untrained girls.

**Table 5.**
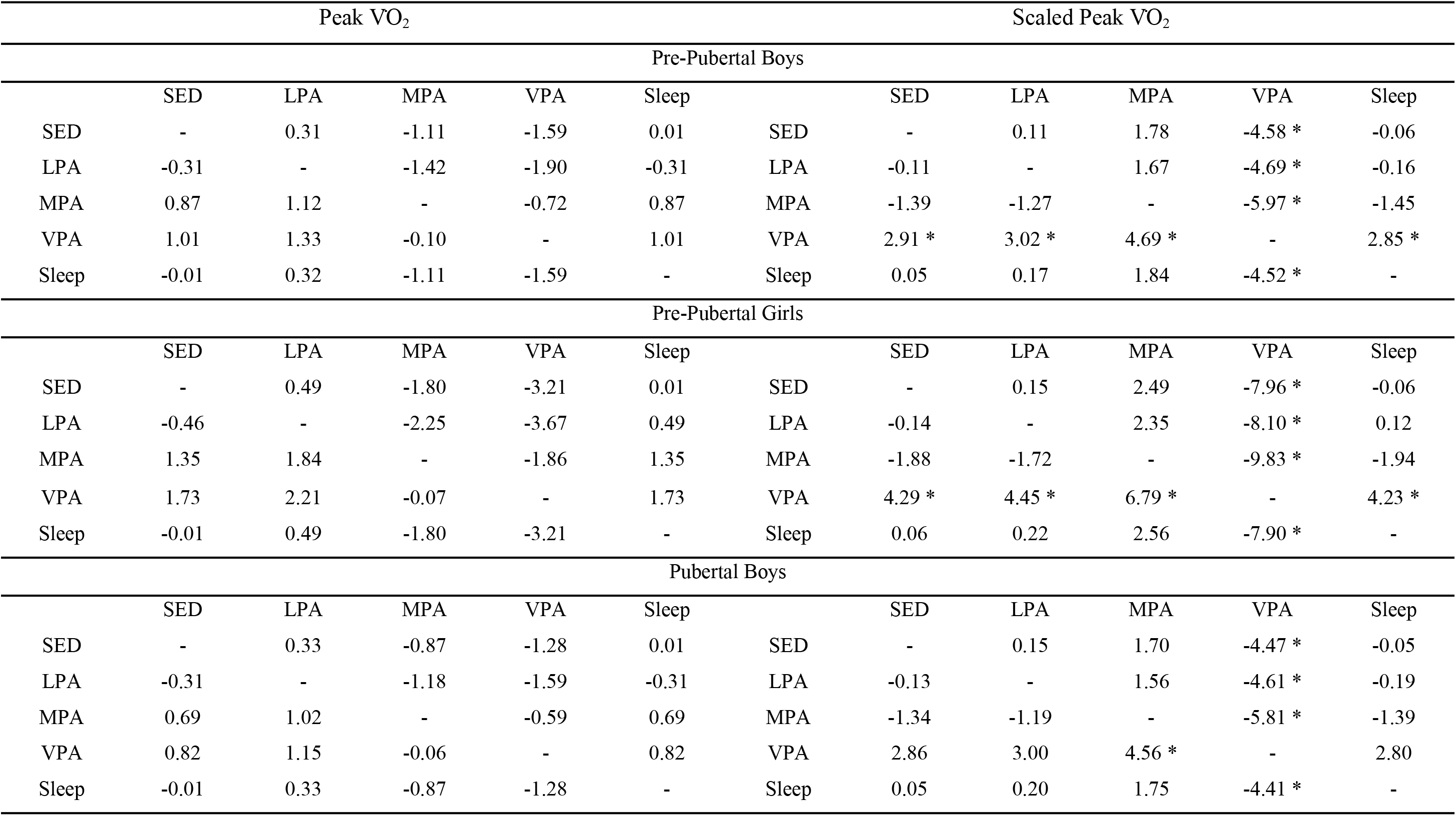
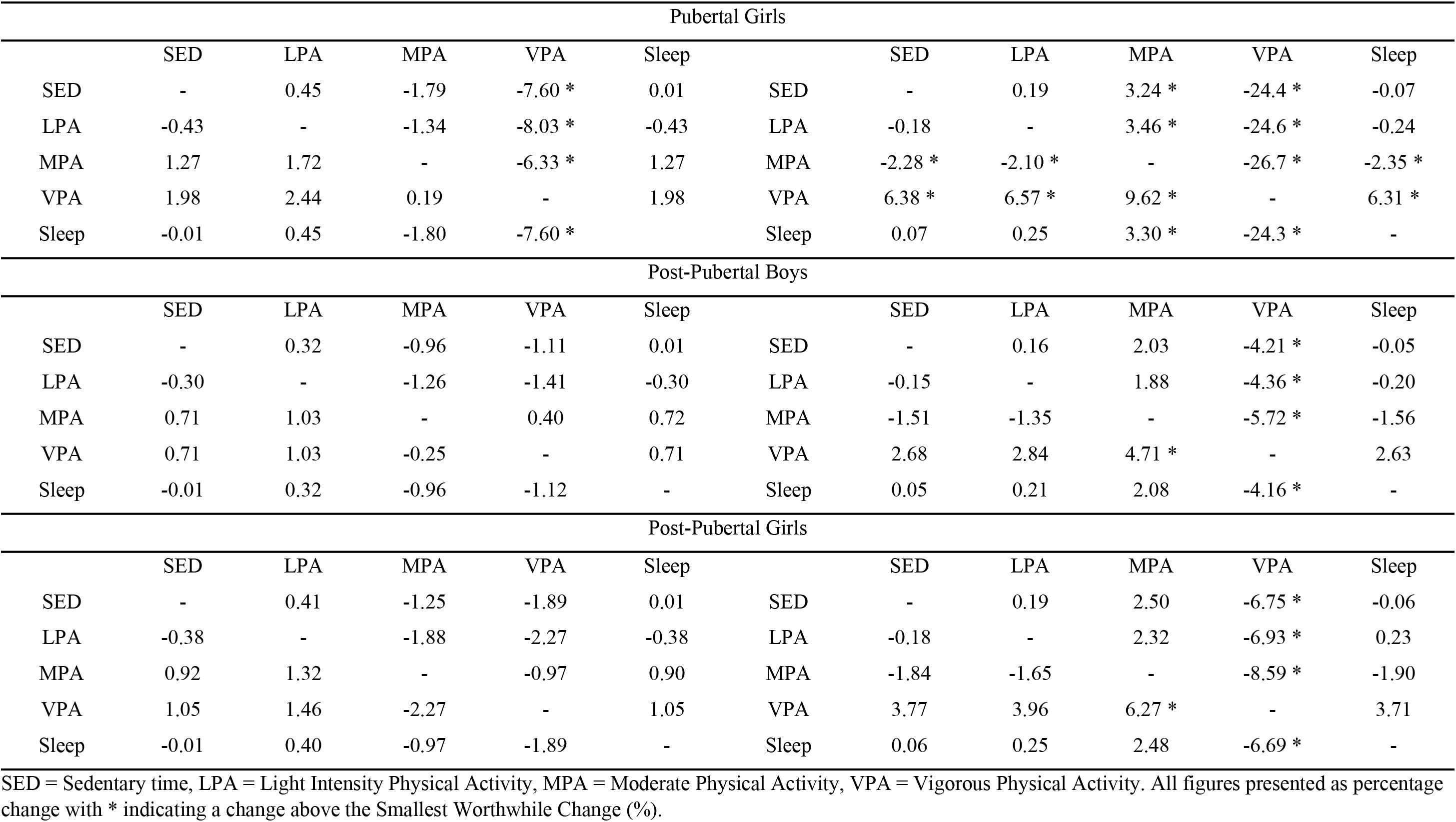
Change matrices of reallocating 10 minutes from the behaviour in columns to the behaviour in the rows on peak VO_2_ (l·min^-1^) and scaled peak VO_2_ (ml-kg^-b^·min^-1^) in trained children and adolescents, presented as percentage change.

| Peak VO<sub>2</sub> |  |  |  |  |  | Scaled Peak VO<sub>2</sub> |  |  |  |  |  |
| --- | --- | --- | --- | --- | --- | --- | --- | --- | --- | --- | --- |
| Pre-Pubertal Boys |  |  |  |  |  |  |  |  |  |  |  |
|  | SED | LPA | MPA | VPA | Sleep |  | SED | LPA | MPA | VPA | Sleep |
| SED | - | 0.31 | -1.11 | -1.59 | 0.01 | SED | - | 0.11 | 1.78 | -4.58 * | -0.06 |
| LPA | -0.31 | - | -1.42 | -1.90 | -0.31 | LPA | -0.11 | - | 1.67 | -4.69 * | -0.16 |
| MPA | 0.87 | 1.12 | - | -0.72 | 0.87 | MPA | -1.39 | -1.27 | - | -5.97 * | -1.45 |
| VPA | 1.01 | 1.33 | -0.10 | - | 1.01 | VPA | 2.91 * | 3.02 * | 4.69 * | - | 2.85 * |
| Sleep | -0.01 | 0.32 | -1.11 | -1.59 | - | Sleep | 0.05 | 0.17 | 1.84 | -4.52 * | - |
| Pre-Pubertal Girls |  |  |  |  |  |  |  |  |  |  |  |
|  | SED | LPA | MPA | VPA | Sleep |  | SED | LPA | MPA | VPA | Sleep |
| SED | - | 0.49 | -1.80 | -3.21 | 0.01 | SED | - | 0.15 | 2.49 | -7.96 * | -0.06 |
| LPA | -0.46 | - | -2.25 | -3.67 | 0.49 | LPA | -0.14 | - | 2.35 | -8.10 * | 0.12 |
| MPA | 1.35 | 1.84 | - | -1.86 | 1.35 | MPA | -1.88 | -1.72 | - | -9.83 * | -1.94 |
| VPA | 1.73 | 2.21 | -0.07 | - | 1.73 | VPA | 4.29 * | 4.45 * | 6.79 * | - | 4.23 * |
| Sleep | -0.01 | 0.49 | -1.80 | -3.21 | - | Sleep | 0.06 | 0.22 | 2.56 | -7.90 * | - |
| Pubertal Boys |  |  |  |  |  |  |  |  |  |  |  |
|  | SED | LPA | MPA | VPA | Sleep |  | SED | LPA | MPA | VPA | Sleep |
| SED | - | 0.33 | -0.87 | -1.28 | 0.01 | SED | - | 0.15 | 1.70 | -4.47 * | -0.05 |
| LPA | -0.31 | - | -1.18 | -1.59 | -0.31 | LPA | -0.13 | - | 1.56 | -4.61 * | -0.19 |
| MPA | 0.69 | 1.02 | - | -0.59 | 0.69 | MPA | -1.34 | -1.19 | - | -5.81 * | -1.39 |
| VPA | 0.82 | 1.15 | -0.06 | - | 0.82 | VPA | 2.86 | 3.00 | 4.56 * | - | 2.80 |
| Sleep | -0.01 | 0.33 | -0.87 | -1.28 | - | Sleep | 0.05 | 0.20 | 1.75 | -4.41 * | - |

| Pubertal Girls |  |  |  |  |  |  |  |  |  |  |  |
| --- | --- | --- | --- | --- | --- | --- | --- | --- | --- | --- | --- |
|  | SED | LPA | MPA | VPA | Sleep |  | SED | LPA | MPA | VPA | Sleep |
| SED | - | 0.45 | -1.79 | -7.60 * | 0.01 | SED | - | 0.19 | 3.24 * | -24.4 * | -0.07 |
| LPA | -0.43 | - | -1.34 | -8.03 * | -0.43 | LPA | -0.18 | - | 3.46 * | -24.6 * | -0.24 |
| MPA | 1.27 | 1.72 | - | -6.33 * | 1.27 | MPA | -2.28 * | -2.10 * | - | -26.7 * | -2.35 * |
| VPA | 1.98 | 2.44 | 0.19 | - | 1.98 | VPA | 6.38 * | 6.57 * | 9.62 * | - | 6.31 * |
| Sleep | -0.01 | 0.45 | -1.80 | -7.60 * |  | Sleep | 0.07 | 0.25 | 3.30 * | -24.3 * | - |
| Post-Pubertal Boys |  |  |  |  |  |  |  |  |  |  |  |
|  | SED | LPA | MPA | VPA | Sleep |  | SED | LPA | MPA | VPA | Sleep |
| SED | - | 0.32 | -0.96 | -1.11 | 0.01 | SED | - | 0.16 | 2.03 | -4.21 * | -0.05 |
| LPA | -0.30 | - | -1.26 | -1.41 | -0.30 | LPA | -0.15 | - | 1.88 | -4.36 * | -0.20 |
| MPA | 0.71 | 1.03 | - | 0.40 | 0.72 | MPA | -1.51 | -1.35 | - | -5.72 * | -1.56 |
| VPA | 0.71 | 1.03 | -0.25 | - | 0.71 | VPA | 2.68 | 2.84 | 4.71 * | - | 2.63 |
| Sleep | -0.01 | 0.32 | -0.96 | -1.12 | - | Sleep | 0.05 | 0.21 | 2.08 | -4.16 * | - |
| Post-Pubertal Girls |  |  |  |  |  |  |  |  |  |  |  |
|  | SED | LPA | MPA | VPA | Sleep |  | SED | LPA | MPA | VPA | Sleep |
| SED | - | 0.41 | -1.25 | -1.89 | 0.01 | SED | - | 0.19 | 2.50 | -6.75 * | -0.06 |
| LPA | -0.38 | - | -1.88 | -2.27 | -0.38 | LPA | -0.18 | - | 2.32 | -6.93 * | 0.23 |
| MPA | 0.92 | 1.32 | - | -0.97 | 0.90 | MPA | -1.84 | -1.65 | - | -8.59 * | -1.90 |
| VPA | 1.05 | 1.46 | -2.27 | - | 1.05 | VPA | 3.77 | 3.96 | 6.27 * | - | 3.71 |
| Sleep | -0.01 | 0.40 | -0.97 | -1.89 | - | Sleep | 0.06 | 0.25 | 2.48 | -6.69 * | - |

SED = Sedentary time, LPA = Light Intensity Physical Activity, MPA = Moderate Physical Activity, VPA = Vigorous Physical Activity. All figures presented as percentage change with \* indicating a change above the Smallest Worthwhile Change (%).

**Table 6.** Change matrices of reallocating 10 minutes from the behaviour in columns to the behaviour in the rows on peak VO_2_ (l·min^-1^) and scaled peak VO_2_ (ml·kg^-b^·min^-1^) in untrained children and adolescents, presented as percentage change.

| Peak VO<sub>2</sub> |  |  |  |  |  | Scaled Peak VO<sub>2</sub> |  |  |  |  |  |
| --- | --- | --- | --- | --- | --- | --- | --- | --- | --- | --- | --- |
| Pre-Pubertal Boys |  |  |  |  |  |  |  |  |  |  |  |
|  | SED | LPA | MPA | VPA | Sleep |  | SED | LPA | MPA | VPA | Sleep |
| SED | - | 0.63 | -1.34 | -3.17 * | 0.01 | SED | - | 0.23 | 2.11 | -8.93 * | -0.07 |
| LPA | -0.59 | - | -1.93 | -3.75 * | -0.59 | LPA | -0.22 | - | 1.90 | -9.14 * | -0.28 |
| MPA | 1.05 | 1.69 | - | -2.11 | 1.05 | MPA | -1.66 | -1.42 | - | -10.59 * | -1.72 |
| VPA | 1.62 | 2.25 | 0.27 | - | 1.62 | VPA | 4.58 | 4.81 | 6.69 * | - | 4.51 |
| Sleep | -0.01 | 0.63 | -1.34 | -3.17 |  | Sleep | 0.07 | 0.30 | 2.18 | -9.15 * | - |
| Pre-Pubertal Girls |  |  |  |  |  |  |  |  |  |  |  |
|  | SED | LPA | MPA | VPA | Sleep |  | SED | LPA | MPA | VPA | Sleep |
| SED | - | 2.72 | -3.93 | -8.31 * | 0.01 | SED | - | 0.88 | 5.17 * | -19.43 * | -0.09 |
| LPA | -2.21 | - | -6.14 * | -10.52 * | -2.20 | LPA | -0.71 | - | 4.46 * | -20.13 * | -0.79 |
| MPA | 2.48 | 5.20 * | - | -5.84 * | 2.48 | MPA | -3.25 | -2.36 | - | -22.67 * | -3.33 |
| VPA | 2.86 | 5.58 * | -1.07 | - | 2.86 | VPA | 6.70 * | 7.57 * | 11.86 * | - | 6.61 * |
| Sleep | -0.01 | 2.72 | -3.94 | -8.31 * |  | Sleep | 0.06 | 0.96 | 5.24 * | -19.34 * | - |
| Pubertal Boys |  |  |  |  |  |  |  |  |  |  |  |
|  | SED | LPA | MPA | VPA | Sleep |  | SED | LPA | MPA | VPA | Sleep |
| SED | - | 0.53 | -1.12 | -3.00 | 0.01 | SED | - | 0.27 | 2.22 | -10.64 * | -0.07 |
| LPA | -0.53 | - | -1.64 | -3.53 | -0.53 | LPA | -0.25 | - | 1.97 | -10.89 * | -0.32 |
| MPA | 0.87 | 1.44 | - | -2.13 | 0.87 | MPA | -1.72 | -1.45 | - | -12.36 * | -1.79 |
| VPA | 1.40 | 1.97 | 0.28 | - | 1.40 | VPA | 4.96 * | 5.23 * | 7.18 * | - | 4.89 * |
| Sleep | -0.01 | 0.57 | -1.12 | -3.01 | - | Sleep | 0.07 | 0.34 | 2.29 | -10.58 * | - |

| Pubertal Girls |  |  |  |  |  |  |  |  |  |  |  |
| --- | --- | --- | --- | --- | --- | --- | --- | --- | --- | --- | --- |
|  | SED | LPA | MPA | VPA | Sleep |  | SED | LPA | MPA | VPA | Sleep |
| SED | - | 0.77 | -1.43 | -3.85 | 0.01 | SED | - | 0.33 | 2.58 | -12.32 * | -0.08 |
| LPA | -0.71 | - | -2.15 | -4.57 | -0.71 | LPA | -0.30 | - | 2.28 | -12.63 * | -0.38 |
| MPA | 1.13 | 1.90 | - | -2.72 | 1.13 | MPA | -2.02 | -1.68 | - | -14.34 * | -2.10 |
| VPA | 1.84 | 2.62 | 2.72 | - | 1.84 | VPA | 5.90 * | 6.22 * | 4.90 * | - | 5.81 * |
| Sleep | -0.01 | 0.77 | -1.44 | -3.86 |  | Sleep | 0.08 | 0.41 | 2.66 | -12.25 * | - |
| Post-Pubertal Boys |  |  |  |  |  |  |  |  |  |  |  |
|  | SED | LPA | MPA | VPA | Sleep |  | SED | LPA | MPA | VPA | Sleep |
| SED | - | 0.62 | -1.23 | -2.30 | 0.01 | SED | - | 0.32 | 2.67 | -8.89 * | -0.06 |
| LPA | -0.56 | - | -1.79 | -2.86 | -0.56 | LPA | -0.29 | - | 2.38 | -9.18 * | -0.35 |
| MPA | 0.89 | 1.51 | - | -1.41 | 0.89 | MPA | -1.93 | -1.60 | - | -10.82 * | -1.99 |
| VPA | 1.12 | 1.74 | -0.11 | - | 1.12 | VPA | 4.34 * | 4.66 * | 7.00 * | - | 4.27 * |
| Sleep | -0.01 | 0.62 | -1.23 | -2.30 | - | Sleep | 0.06 | 0.38 | 2.73 * | -8.83 * | - |
| Post-Pubertal Girls |  |  |  |  |  |  |  |  |  |  |  |
|  | SED | LPA | MPA | VPA | Sleep |  | SED | LPA | MPA | VPA | Sleep |
| SED | - | 0.44 | -1.54 | -3.55 | 0.01 | SED | - | 0.02 | 3.10 | -12.70 * | -0.08 |
| LPA | -0.42 | - | -1.96 | -3.96 | -0.41 | LPA | -0.19 | - | 2.91 | -12.89 * | -0.27 |
| MPA | 1.14 | 1.57 | - | -2.41 | 1.14 | MPA | -2.28 | -2.07 | - | -14.98 * | -2.36 |
| VPA | 1.59 | 2.02 | 1.59 | - | 1.59 | VPA | 5.70 * | 5.91 * | 5.68 * | - | 5.63 * |
| Sleep | -0.01 | 0.44 | -1.54 | -3.54 | - | Sleep | 0.08 | 0.28 | 3.17 | -12.62 * | - |

SED = Sedentary time, LPA = Light Physical Activity, MPA = Moderate Physical Activity, VPA = Vigorous Physical Activity. All figures presented as percentage change with \* indicating a change greater than the Smallest Worthwhile Change (%).

Allometrically scaled VO_2max_ significantly decreased, irrespective of sex, maturity, or training status, when 10 minutes of VPA was reallocated to any other behaviour (Tables 5 and 6). Moreover, allometrically scaled VO_2max_ tended to increase when time spent in other movement behaviours was reallocated to VPA. In isolation, both SED and sleep were not significant predictors of absolute, or scaled, VO_2max_ (Table 4) however, displacing SED with any other behaviour had a negligible effect on VO_2max_ (0.01 – 1.98%; Tables 5 and 6) unless it was displaced with VPA where it was associated with an increase of 2.68 – 6.38% in scaled VO_2max_ only. Similarly, compositions with 10 minutes less sleep, and 10 minutes more VPA, were associated with a 2.85 – 6.31% greater allometrically scaled VO_2max_ irrespective of sex, training, and maturation. The effect of reallocating time to LPA, SED, or sleep on VO_2max_ was negligible, irrespective of how VO_2max_ was expressed, sex, maturity, or training status.

## 4. Discussion

This is the first study to examine the joint effects of various movement behaviours (SED, LPA, MPA, VPA and Sleep), using a five-part compositional analysis, on absolute and allometrically scaled VO_2max_ in trained and untrained children and adolescents. The main findings of the present study were that allocating time to, and removing time from, VPA significantly increased and decreased allometrically scaled VO_2max_, respectively, regardless of sex, training, or maturity status. Moreover, this study suggests that VPA is potentially 2.4 – 4.7% more potent in eliciting an improvement in VO_2max_ over a 10-minute period in children and adolescents. These findings therefore highlight that intensity of PA may be of paramount importance in determining peak VO_2_, especially in girls.

Engaging in 10 minutes more VPA, irrespective of which behaviour it displaces, significantly increases both absolute and allometrically scaled VO_2max_, regardless of training status. Moreover, of importance, untrained children were predicted to have a larger magnitude of change for the same 10-minute reallocation, in accord with the review of McNarry & Jones [8] which concluded that baseline fitness significantly impacts the magnitude of change experienced to a given stimulus. More specifically, Mahon [37] reported that 52% of the inter-individual variation in participants responses to a training stimulus can be explained by baseline VO_2max_. The findings of the current study are, however, discordant with Carson et al. [16] who reported no significant differences when reallocating time to, or from, any movement behaviour. Such discrepancies may be explained by the use of a proxy measure of VO_2max_ and not accounting for maturation or training status in the earlier study, which are critical when assessing cardiorespiratory fitness in children and adolescents [10,11,27].

The present study supports the notion that children and adolescents may require a vigorous stimulus to significantly improve absolute and scaled VO_2max_ [8,10,12–14]. Moreover, increasing levels of VPA is more time efficient, a commonly cited barrier to physical activity [38], than MPA thus more targeted public health messaging may be needed. However, of concern however, the current findings suggest that children and adolescents may need to increase their time spent in VPA by over 50%. Indeed, the average time spent in VPA in this study was 17.1 ± 12.7 mins·day^-1^, which is in accord with the levels reported in the millennium cohort study (19.9 ± 10.6 mins·day^-1^). Considering the limited success at increasing PA in many interventions to date [38,39], and the small magnitude of increases in VPA reported even in those considered successful [40], the current findings highlight the need to drastically change our approach to PA promotion. Indeed, these findings could be speculated to support the contention suggested by many authors that HIIT may represent an important public health intervention tool [29,41].

Dencker et al. [12] reported weak, but significant, correlations between VPA and peak VO_2_ (r^2^ = 0.32) and allometrically scaled peak VO_2_ (r^2^ = 0.27). Furthermore, the most recent review of the relationship between PA and peak VO_2_ in youth concluded that, despite decades of research, there was still no consensus [11]. These equivocal findings may be related to the reliance on techniques that fail to account for the inter-related and inherently constrained nature of PA behaviours, leading to spurious conclusions [18,25,26]. Moreover, the reliance on ratio scaling VO_2max_ potentially creates spurious associations [15,42]. Of note, when VO_2max_ was allometrically scaled by body mass, the overall PA composition explained ~11% less variance compared to absolute VO_2max_. This may be due, at least in part, to physically active children having a higher lean body mass (LBM) than their sedentary counterparts [43], indicating that differences in body composition may also be critical when determining the effect of re-allocating PA. Nevertheless, the PA composition still explained ~37.7% of the variance in allometrically scaled VO_2max_, demonstrating the powerful influence of habitual PA on aerobic fitness.

The finding that allocating time to MPA decreased allometrically scaled peak VO_2_ was surprising. These associations could be due at least in part to the high amount of VPA undertaken by the trained group within this study and consequently if time is removed, and replaced with MPA, it will have a negative impact on peak VO_2_. Future research is required to explore the interaction of MPA and VPA in other athletic populations to confirm this hypothesis. Of note, the fixed time reallocation used within compositional analysis studies to date [18,20], and in the present study, may over-estimate the magnitude of change in a given variable. More specifically, a 10-minute change in VPA constituted a ~50% increase in VPA within the current sample but the same 10-minute re-allocation only initiated a 1.9% increase in SED time. Therefore, a greater insight into the independent, and interactive, effects of movement behaviours on peak VO_2_ may be gained by investigating the effects of the same percentage change in movement behaviours on peak VO_2_. Nevertheless, evidence is emerging that the intensity of PA may be critical in improving both performance and health-related parameters in paediatric populations [18,20,25] and thus VPA should be encouraged, as opposed to MPA, to engender the greatest long-term health benefits.

Sedentary time in isolation was not a significant predictor of both absolute, and allometrically, scaled VO_2max_ in children and adolescents within the current study irrespective of sex, maturation, and training status. This is in direct conflict with a growing body of literature suggesting that excess sedentary time could lead to a decreased VO_2max_ [21,22] and suggests instead of SED being the problem *per se* it is potentially the PA behaviour it replaces. More specifically, significant increases in VO_2max_ were only associated with compositions where the amount of time spent sedentary decreased and was replaced with VPA. However, whilst displacing SED with LPA and MPA was also associated with an increased VO_2max_ if these behavioural changes were introduced as part of a wider health initiative, they could still contribute to improving the health of the nation. As we move, and transition, out of the COVID-19 pandemic and as restrictions continue to ease public health messaging should highlight the need to avoid displacing PA with SED, put particularly VPA.

Future research should seek to implement targeted interventions informed by compositional analyses, to ascertain the required duration needed to elicit the changes predicted. This is of particular importance as a plethora of research has investigated the influence of different training methodologies on both absolute and allometrically scaled VO_2max_, with their effectiveness being reviewed elsewhere [8,10]. One major issue with the majority of paediatric training studies to date is the lack of accounting for changes in habitual PA levels across the intervention period, and this could help explain the equivocal findings of some intervention types [10,43]. Specifically, most interventions elicit an improvement in VO_2max_ of 5-6% increasing to ~10.4% when allometrically scaled [10]. Thus, if the predicted increases in absolute, scaled, VO_2max_ can be achieved over 4 – 6 weeks, the typical length of most training interventions [8–11], then it is not possible to delineate whether the improvements in VO_2max_ are training-related or stem from increases in habitual PA.

Whilst there are numerous strengths associated with this study, such as the use of a novel five-part compositional analysis approach, allometrically scaling peak VO_2_, and accounting for training, maturity and sex differences, there are limitations which must be acknowledged. Firstly, a relatively low wear-time criteria was set of any three days with at least eight hours of wear-time; a more stringent wear-time criteria could potentially influence the relationships established between PA metrics and peak VO_2_. Nevertheless, this wear-time has been validated in a paediatric population [34] and was used to maximise participant inclusion within the study. Secondly, the cross-sectional study does not allow the duration over which the habitual changes need to be maintained for to observe the associated changes. Thirdly, the small sample size compared to other studies of this type [18,25] and the representativity of this population needs to be considered when interpreting the results of this study. Finally, the applicability of cycle derived peak VO_2_ to habitual PA levels is contentious, and therefore future research should endeavour to establish peak VO_2_ using treadmills to maximise specificity, and to establish whether these findings persist.

## 5. Conclusion

In conclusion, the proportion of time VPA is a significant predictor of allometrically scaled peak VO_2_ in children and adolescents, independent of training, sex, and maturity status even when the proportion of time spent in other behaviours is considered. Moreover, reallocating time from VPA in pre-pubertal children significantly predicts a reduced absolute peak VO_2_, potentially highlighting the importance of promoting VPA in pre-pubertal children. Future research should seek to establish the duration of targeted PA interventions needed to elicit the significant changes predicted from compositional analyses and report the individual levels of MPA and VPA to ascertain the relative importance of VPA for current, and future, health in children and adolescents.

## Acknowledgements

The authors would like to thank Dr Rachel Hughes and Tim Evans for their help with the design and recruitment for the study. The authors would also like to thank the children and young people and their parents / guardians for taking their time to complete this study.

## Supplementary Information

**S1 Table 1. Smallest worthwhile change in all sub-groups for peak VO2 (l·min-1) and allometrically scaled peak VO2 (ml·kg-b·min-1).** SWC = Smallest Worthwhile change, SWC (%) = Smallest worthwhile change as a percentage of the individual group mean.

**S1 Figure 1. Ternary heat plots for 24-hour movement behaviours and predicted VO_2max_.** Ternary heat plots of all PA behaviours with expected peak VO2 values for all sub-groups with a) trained athletes; b) untrained controls; c) all boys; d) all girls; e) pre-pubertal children; f) pubertal adolescents; and g) post-pubertal adolescents,

**S1 Figure 2. Ternary heat plots for 24-hour movement behaviours and predicted allometrically scaled VO_2max_.** Ternary plots of all PA behaviours with expected scaled peak VO2 values for all sub-groups with a) trained athletes; b) untrained controls; c) all boys; d) all girls; e) pre-pubertal children; f) pubertal adolescents; and g) post-pubertal adolescent.

